# CD8scape: an accessible, command-line tool for predicting viral escape from the CD8+ T cell response

**DOI:** 10.64898/2026.04.20.719634

**Authors:** Ewan W Smith, Joseph Hughes, David L Robertson, Christopher J R Illingworth

## Abstract

The CD8^+^ T cell response is a critical component of antiviral immunity, particularly in hosts who are immunocompromised or undergoing B cell-depleting therapy, such as rituximab. As viral evolution can lead to escape from CD8^+^ T cell recognition, tools that predict such escape are increasingly relevant. Here, we present CD8scape, an accessible command-line tool designed to predict viral escape from the CD8^+^ T cell response based on within-host sequence variation and HLA class I genotype. CD8scape is primarily a Julia wrapper for NetMHCpan v4.2, a neural network–based predictor trained on mass spectrometry–derived peptide presentation data. CD8scape integrates variant data and viral reading frames to identify all overlapping 8–11mer peptides at variant sites in both ancestral and derived states. These peptides are evaluated using NetMHCpan, which outputs eluted ligand (EL) scores as allele-specific percentile ranks to account for differences in MHC binding fastidiousness, and these are passed back to CD8scape itself. For each variant, the best-ranking peptide across all alleles is identified, and a harmonic mean is used to summarize presentation likelihood across the host’s HLA genotype. A fold-change between ancestral and derived harmonic means quantifies the likelihood of immune escape, with values >1 indicating reduced predicted presentation, and therefore a potential escape from the CD8+ T cell response. This is converted to a log_2_ value of this fold-change so that the metric is symmetric around 0, with positive values representing predicted escape. CD8scape can operate with known HLA genotypes or a representative HLA supertype panel for generalizable predictions. We demonstrate our method by application to within-host SARS-CoV-2 evolution in a rituximab-treated patient and discuss its implications for population-level CD8^+^ T cell escape.

## Introduction

CD8+ T cells, otherwise known as “Killer” T cells, are an important component of the host immune response against viral infection. These recognise their antigen, usually an 8-11mer peptide sequence (Townsend and Bodmer, 1989; Silver, Parker and Wiley, 1991), in the context of Major Histocompatibility Complex class I (MHCI) molecules displayed on the surface of host cells (Figure 1A). MHCI is expressed by all nucleated cells and is the central molecule of the immunological self, as well as the platform through which the CD8+ T cell compartment detects and responds to viral infection and cancer (R. M. Zinkernagel and Doherty, 1974; Rolf M. Zinkernagel and Doherty, 1974; Neefjes *et al*., 2011). The genes that encode a human host’s MHCI proteins are held at the Human Leukocyte Antigen (HLA) locus, which are extremely diverse between individuals and across populations, which is the case for the equivalent locus in all mammals. Classical MHCI molecules are encoded by the HLA-A, HLA-B, and HLA-C genes, which allow a single human host to express up to six different isoforms of MHCI (Dendrou *et al*., 2018). In response to antigen recognition, the CD8+ T cell releases stored granules which contain factors which cause the controlled death of the target cell (Figure 1B) (Peters *et al*., 1991). One of these is perforin, a protein that facilitates the entry of other cytotoxic molecules into the target cell by creating transmembrane pores. Others are granzymes, serine proteases with diverse pro-apoptotic functions, including granzyme B, which activates caspases by cleavage leading to the cleavage of diverse downstream substrates and the permeabilization of the mitochondrial outer membrane; granzyme A, which induces a caspase-independent cell death involving single-stranded DNA damage and disruption of mitochondrion function; and others, including granzymes C, K, and M, which show varying roles and substrate targets (Podack and Dennert, 1983; Pinkoski *et al*., 1998; Martinvalet, Zhu and Lieberman, 2005; Podack and Munson, 2016; Hay and Slansky, 2022).

**Figure 1.**
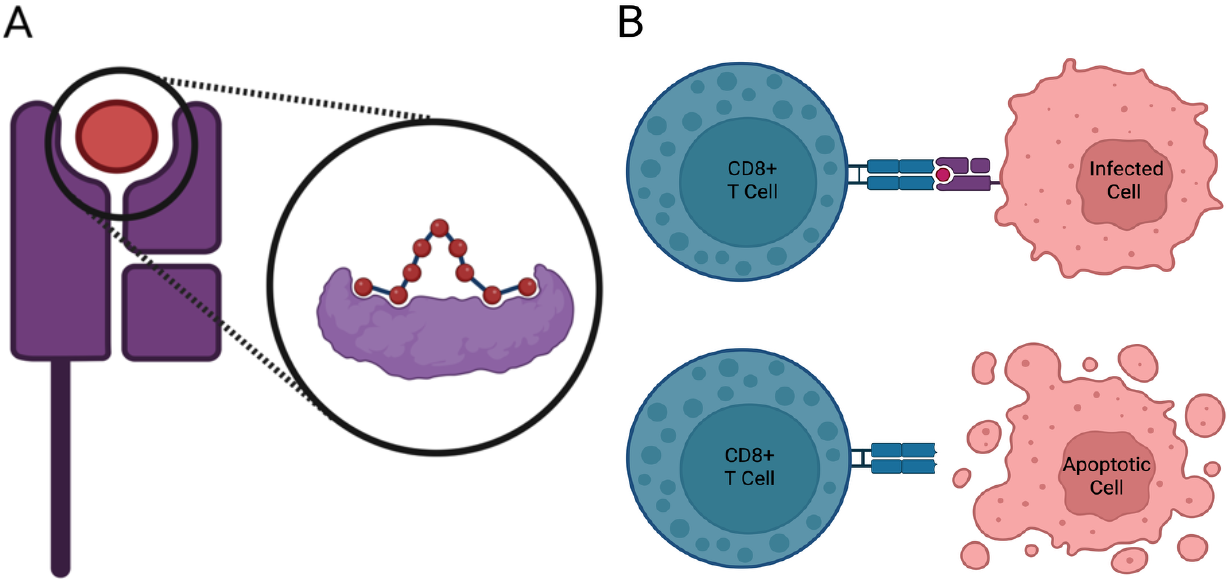
Mechanism of CD8+ T cell recognition. **A)** A peptide antigen (red) presented in the binding cleft of a Class I MHC molecule (purple). **B)** Recognition of the peptide– MHC complex by an activated CD8+ T cell (blue), triggering destruction of the presenting cell (pink). Created in BioRender.com.

The CD8+ T cell compartment forms a core part of the host immune response against viral infection. For example, CD8+ T cells play a critical role in the immune response to HIV (Walker and McMichael, 2012). While discussion of the evolution of respiratory viruses, and in particular attempts to predict virus evolution, have tended to focus upon the role of B-cell mediated antibody responses (Petrova and Russell, 2018), T cell responses also play a critical role in host defence (Jansen *et al*., 2019; Bertoletti, Tan and Le Bert, 2021; De *et al*., 2023). T cell responses are of particular importance in hosts who are immunocompromised in such a way that leaves them without a normally functioning B cell compartment. Rituximab, a chimeric monoclonal antibody targets CD20, a B cell marker, and depletes these cells. CD20 is present across the B cell lineage from the pre-B stage but is not present on terminally differentiated plasma cells. The immunological outcome of this pattern of CD20 expression is that a host receiving rituximab therapy or another anti-CD20 monoclonal antibody therapy is likely to maintain pre-existing antibody levels in serum, so immediate opportunistic infections are unlikely, but depletion of naÏve and memory B cells leaves the host unable to generate antibody responses against novel antigens (Van Der Kolk *et al*., 2002). This depletion of the B cell compartment outlasts anti-CD20 treatment itself, and for example has been demonstrated to suppress responses to vaccines several months after the completion of therapy (Apostolidis *et al*., 2021).

Rituximab was licensed in the late 1990s for the treatment of relapsed or refractory B cell lymphomas and almost immediately changed the standard of care. It soon became an important part of chemotherapy regimens like R-CHOP, which was shown to be more effective in the treatment of diffuse large B cell lymphomas than the CHOP regimen alone (Coiffier *et al*., 2002). It is now used for a wider variety of B cell malignancies, such as Chronic Lymphocytic Leukaemia and Waldenström’s macroglobulinemia. Over time, use of rituximab in the treatment of autoimmune disease has become more common – in 2006, use in the treatment of rheumatoid arthritis was approved in the US and EU, to be joined by granulomatosis with polyangiitis and microscopic polyangiitis in 2010. In 2017 and 2018 in the EU and US respectively its use was approved for pemphigus vulgaris. Off-label use in other autoimmune diseases, such as systemic lupus erythematosus, multiple sclerosis, and myasthenia gravis, is also becoming more common over time (Delate *et al*., 2020).

The increasing usage of B cell depletion therapy highlights the importance of understanding viral escape from the CD8+ T cell compartment. The netMHCpan software package is a valuable and frequently-used method for CD8+ epitope prediction (Nilsson *et al*., 2025). Our use of this method identified the potential for further automation and more rapid application to viral populations. We therefore introduce CD8scape, an accessible command-line tool which functions as a wrapper for netMHCpan, facilitating the automated submission of tasks and the collation of outputs. We propose CD8scape as a valuable tool for the prediction of viral CD8+ T cell escape.

## Methods

CD8scape is written largely in Julia v1.11.5 (Bezanson *et al*., 2017), including some scripts in Perl v5.34.1 (Christiansen *et al*., 2012). At its core sits netMHCpan v4.2, which performs the prediction of binding and elution between supplied peptides and given MHC molecules (Reynisson *et al*., 2020; Nilsson *et al*., 2025).

CD8scape reads in a list of variants, specified either as a .vcf or as a .out file from SAMFIRE (Illingworth, 2016). It further reads in a consensus genome, specified as a single-line fasta file, and a list of reading frames in the genome, with reading frames taken either as an output from SAMFIRE or in the standard download format from NCBI Virus (Brister *et al*., 2015). These files provide a context-aware understanding of variation in the viral population.

Given this information, CD8scape generates every peptide of lengths 8, 9, 10, and 11 amino acids from the reference sequence that overlaps the variant locus, identifying 8-mers to 11-mers both with and without the substitution edited in (Figure S1). We refer to these as the ancestral and derived states of each peptide.

Generated peptides are passed to NetMHCpan, a machine learning–based model that predicts peptide:MHCI interactions. The output of this method is referred to as the EL score, which represents the likelihood that a given peptide will be naturally presented by the MHC molecule. In order to account for varying permissiveness across MHCI molecules (Paul *et al*., 2013), this is converted to a percentage rank relative to a large set of random natural presented peptides for a given MHCI allele.

### Measuring MHC escape for a known HLA genotype

We index the k-mers containing an allele using the variable *i*. We denote the rank of k-mer *i* containing the ancestral allele with respect to MHCI molecule *m* as *r*_*a,m,i*_, and likewise the rank of k-mer *i* containing the derived allele as *r*_*d,m,i*_. Ranks are expressed as percentiles, bounded between 0 and 100. A high rank, corresponding to a low rank-number, is obtained by a stronger-binding peptide. Following the convention described in the netMHCpan documentation, we defined cutoffs for strong binding, having a rank number in the range 0 to 0.5%, and weak binding, having a rank number in the range 0.5 to 2%. As such, for each MHC allele, the ranks of the strongest binding peptides in the ancestral and derived states are given by

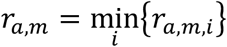

and

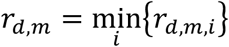

These ranks are processed by CD8scape, producing a measurement for each MHC allele of the extent of escape generated by a specific mutation in the viral genome. Loci for which the best ranks in both the ancestral and derived states are outside of the top 2% are removed from further consideration. We then calculate the log_2_ fold-change:

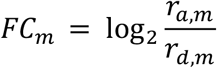

A log_2_ fold-change of greater than 0 indicates a predicted reduction in presentation by allele *m* in the derived state relative to the ancestral state, indicating a potential escape from CD8+T cell recognition.

Optionally, CD8scape also provides a cross-allele measurement of escape, calculating harmonic mean rank values across MHC molecules:

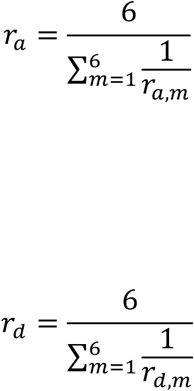

As a combined metric, the harmonic mean highlights peptides that bind strongly to at least one MHC allele, even if their ranks are weaker for others. Here the log_2_ fold-change r_a_/r_d_ between the binding ranks for the ancestral and derived states provides a generic measurement of CD8+ T cell escape.

### Measuring MHC escape where the HLA genotype is unknown

Where the method above considers potential CD8+ T cell escape in the context of a host with a known HLA genotype, an extension of this considered escape for a representative supertype panel of HLA genes, that is a weighted panel of HLA genes that seeks to describe the makeup of the population to which the individual belongs. MHC allele-specific statistics are output for each allele within the population. To calculate a statistic across alleles, we generate a weighted harmonic mean across the different HLA types. Where r_a,m_ is the rank of the strongest binding peptide for HLA type m, we calculate:

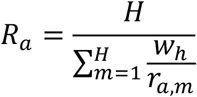

In this manner, CD8scape allows a mean estimate to be made of the extent of escape from the CD8+ T cell compartment for a representative individual in a population. Where a single individual is considered, the strategy of finding the haplotype corresponding to the greatest level of immune escape is a sensible one, given immunodominance. However, in a population supertype panel, where not every individual has every haplotype, the same maximal strategy is not optimal. The harmonic mean provides an approximate measure of cross-population immunity given a supertype panel.

### Simulating MHC escape data

CD8scape also includes the potential to generate a null distribution of escape scores by simulating random mutations across a viral genome. This module first generates all possible nucleic acid substitutions, although options are provided to sample from this by either a count of mutations or by a proportion. A random seed, which can be specified in the settings, ensures the reproducibility of sampled datasets.

Simulated mutations are processed in an identical manner to observed variants to generate an empirical distribution of escape scores that represents the background expectation of mutational impact on antigen presentation.

The simulation framework can be applied either to a known HLA genotype panel for an individual host or to a representative HLA supertype panel. In this way observed variants can be contextualised within both the potential mutational landscape of the host’s known genotype or where that genotype is unknown. This offers an alternative framework for analysing mutation effects. Rather than interpreting escape solely through the fold-change metric, which is highly derived, mutations can be ranked relative to the simulated distribution.

### Assessing collective escape across a variant set

To assess whether a set of observed variants shows collectively elevated escape relative to the simulated null, each observed variant is assigned a percentile within the simulated distribution of log_2_ fold-change scores. Individual percentiles are converted to Z-scores via the probit transform and combined using Stouffer’s method:

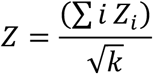

where k is the number of variants with a valid percentile. A one-tailed p-value is derived from the standard normal distribution. As a non-parametric complement, an empirical p-value is computed by drawing an ensemble of random samples of k variants without replacement from the simulated distribution and calculating the mean percentile of each sample. The empirical p-value is the proportion of samples whose mean percentile meets or exceeds the observed mean percentile. Both statistics are reported alongside the per-variant percentiles in the CD8scape output.

## Results

By way of demonstration, we analysed data from the case study of a female Russian patient undergoing chemotherapy that included rituximab for non-Hodgkin’s diffuse B-cell lymphoma and contracted SARS-CoV-2 while in hospital (Stanevich *et al*., 2023). Viral variant data and this patient’s HLA genotype (HLA-A01:01, HLA-A03:01, HLA-B07:02, HLA-B08:01, HLA-C07:01, and HLA-C07:02) were accessed from the supplementary information of this paper. Viral sequence data was accessed from the Sequence Read Archive (accession: PRJNA749008).

28 variants across 12 proteins were found to be non-synonymous and present at a minimum 10% at any time point. One of these, S672^*^ in the RdRp, induces a stop codon and so was removed from further consideration. 7 more variants had both their ancestral and derived state harmonic mean best ranks (HMBRs) above 2, so were removed from further consideration (Figure 2A). 19 variants made it to the end of the analysis pipeline (Figure 2B). In common with the original analysis of these data, the harmonic mean statistic identifies the T504A, T504P, and D821N variants of nsp3 as of potentially high importance in conveying CD8+ T cell escape.

**Figure 2.**
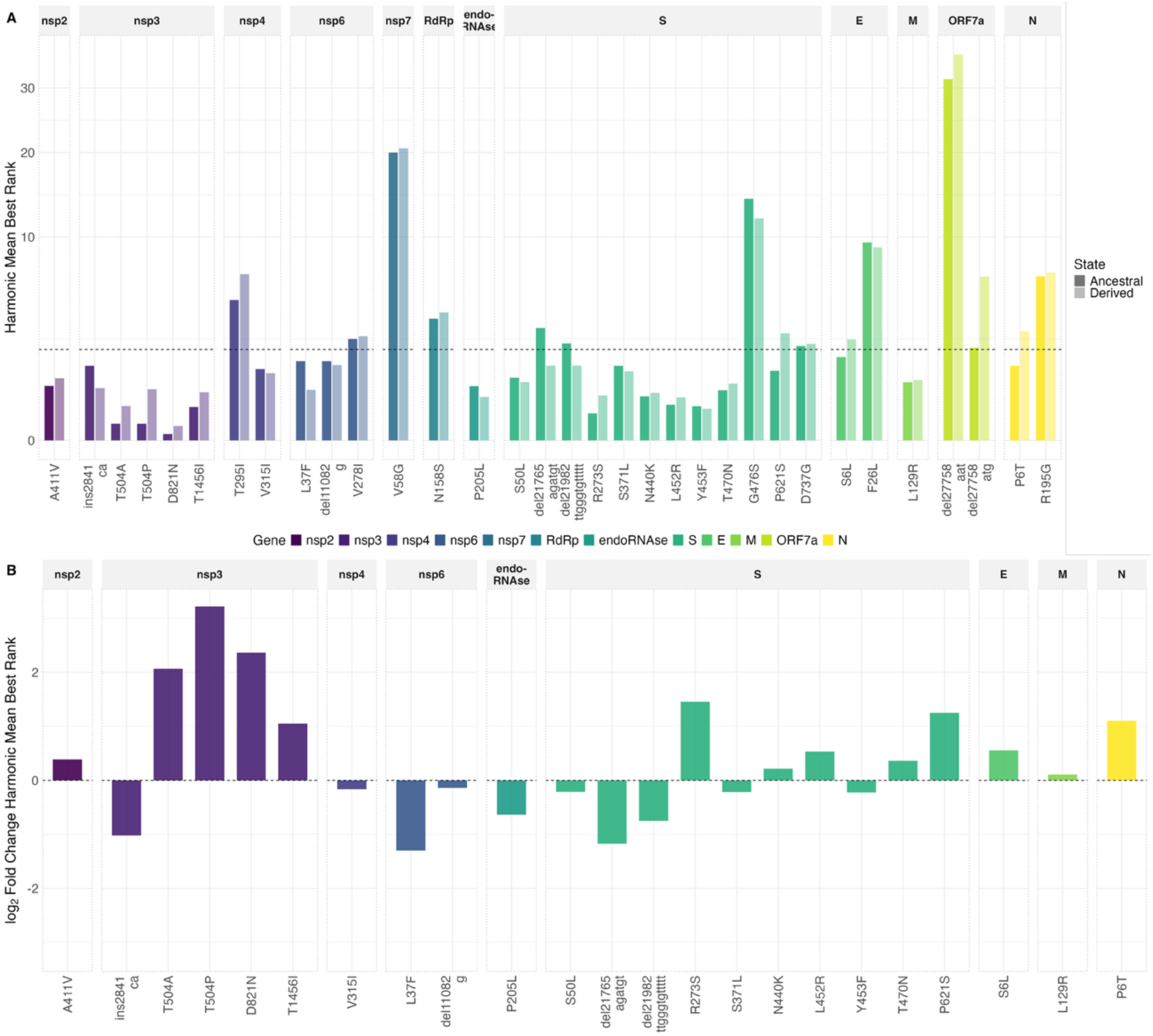
Calculated estimates of CD8+ T cell escape for variants identified in a case of chronic SARS-CoV-2 infection. **A)** Harmonic mean best EL % rank for each variant in the ancestral (darker) and derived (paler) states, shown on a square-root scale. The dashed line marks the 2% binding threshold; variants exceeding this in both states are excluded from further analysis. **B)** Log_2_ fold-change in harmonic mean best EL rank from ancestral to derived state. Positive values indicate predicted weakening of antigen presentation.

Representative supertype frequencies for genes HLA-A, HLA-B, and HLA-C (Figure 3A) were taken from the Moscow Stem Cell Bank entry on the Allele Frequency Net Database (Gonzalez-Galarza *et al*., 2019). The supertype data shows some concordance with those for the patient (Figure 3B). Of the three variants mentioned above, T504A and T504P show evidence of generating potential escape within the local Moscow population as a whole, although the signal for D821N disappears at the population level. One hypothesis in studies of SARS-CoV-2 evolution is that virus evolution in immunocompromised patients contributes more broadly to the evolution of the virus in the global population, with chronic infection generating novel viral variation (Choi *et al*., 2020; Corey and Michael, 2021; Kemp *et al*., 2021). This case provides an insight into how such evolution might occur, with variants identified in a chronically-infected patient having potentially broader population-wide impact.

**Figure 3.**
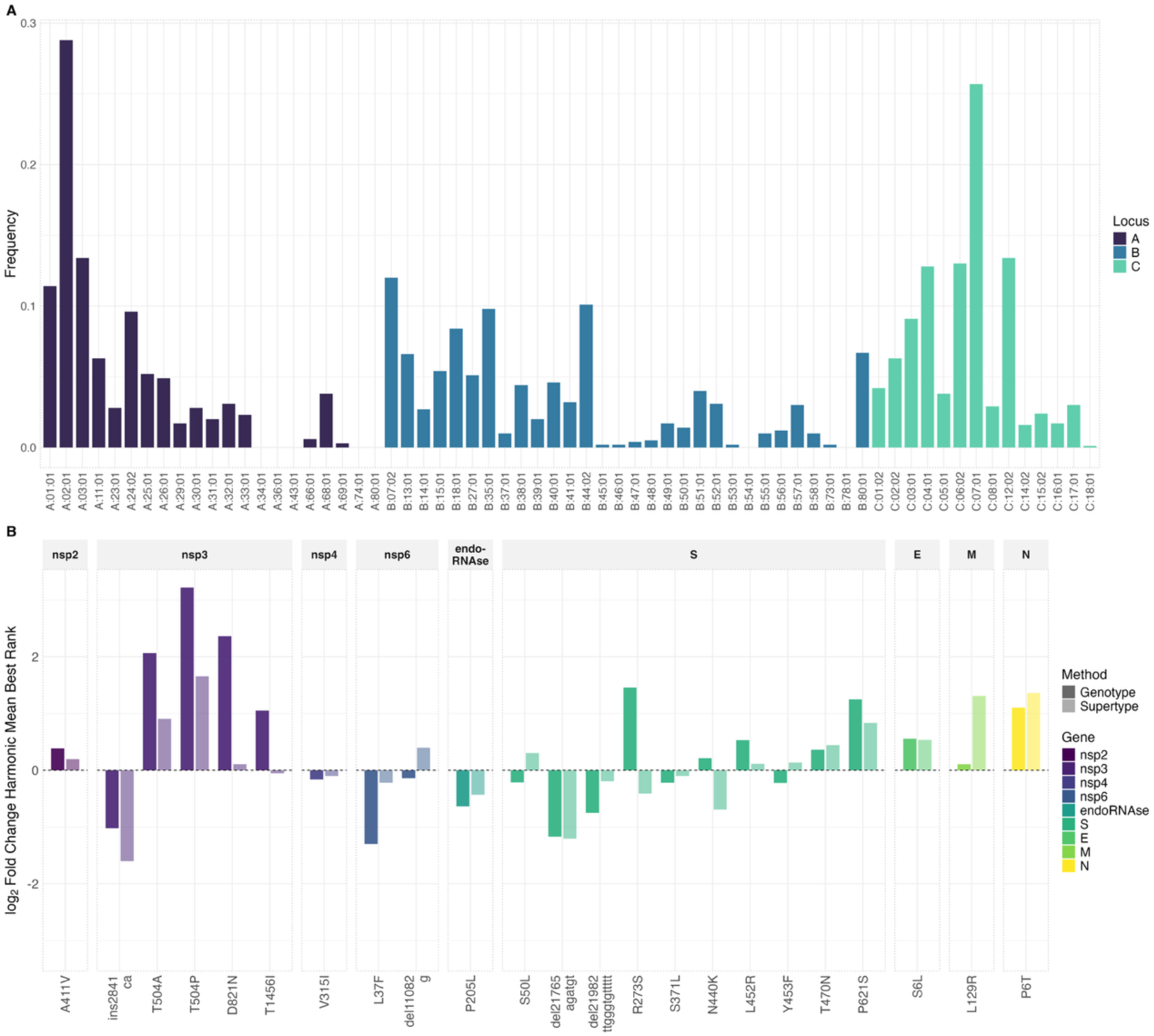
Calculated estimates of CD8+ T cell escape within a local population for variants identified in a case of chronic SARS-CoV-2 infection. **A)** HLA allele frequencies by locus for the representative Moscow supertype panel. **B)** Log_2_ fold-change in harmonic mean best EL rank for each variant, comparing the patient genotype panel (darker) with the representative supertype panel (paler).

Individual output values calculated for each allele in the patient’s genotype suggested underlying mechanisms aligning individual and population-wide effects (Figure 4A). In many cases, the MHC allele generating the strongest signal of escape is either HLA-A01:01 or HLA-B07:02. The former of these is relatively common in the local population, featuring in approximately 10% of individuals. Rank values are shown ordered by allele in Figure 4B.

**Figure 4.**
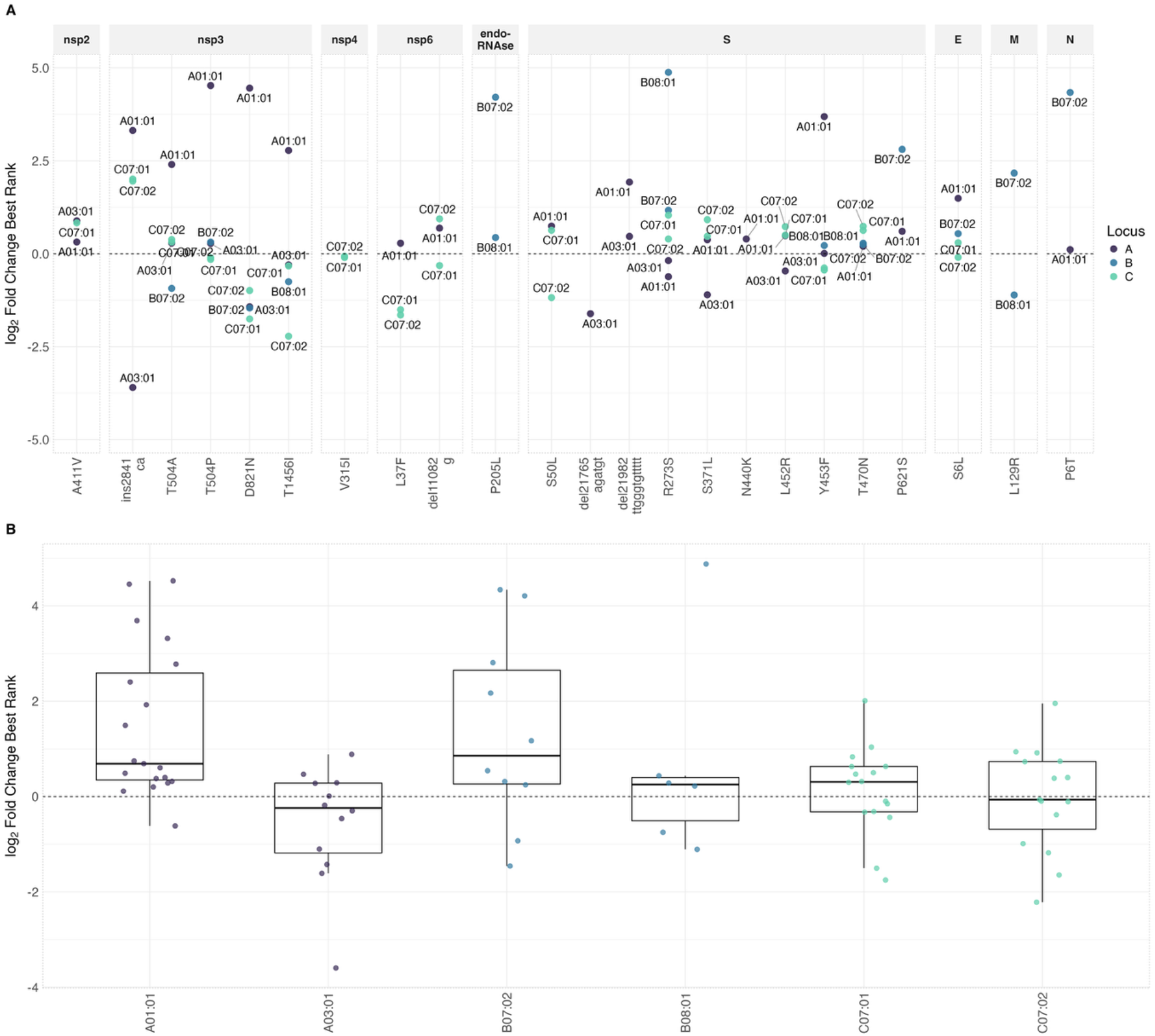
Haplotype-specific estimates of CD8+ T cell escape for variants. **A**) Log_2_ fold-change in best EL rank for each variant–allele combination passing the 2% ancestral binding threshold, coloured by HLA locus. **B)** Distribution of per-allele log_2_ fold-change values across variants, grouped by HLA allele.

Considering the full range of potential non-synonymous mutations in the initial consensus SARS-CoV-2 genome produced a distribution of HMBR scores that was centred roughly on zero (Figure S2A). Comparing calculated scores with the distribution provides an estimate of the significance of the individual HMBR scores of different variants. Assessed on this level, only the T504P and D821N mutations were in the top 5% of variants (Figure S2B).

To assess whether the observed variants collectively show elevated escape relative to the simulated null, individual percentiles were combined using Stouffer’s method and an empirical permutation test. Under the HMBR measure, which requires strong binding in at least one ancestral k-mer across alleles, 23 variants passed filtering. These do not show a statistically significant collective escape signal (Stouffer Z = 1.36, p = 0.087; empirical p = 0.103, Figure 5). When each variant is instead represented by its highest-escaping allele, 28 variants are retained, as the per-allele filtering criterion requires only that a single allele shows strong ancestral binding rather than a cross-allele harmonic mean. Under this measure, a significant collective escape signal emerges (Stouffer Z = 2.38, p = 0.009; empirical p = 0.009, Figure 5), suggesting that escape is concentrated in specific allele–variant pairs rather than distributed broadly across the genotype.

**Figure 5.**
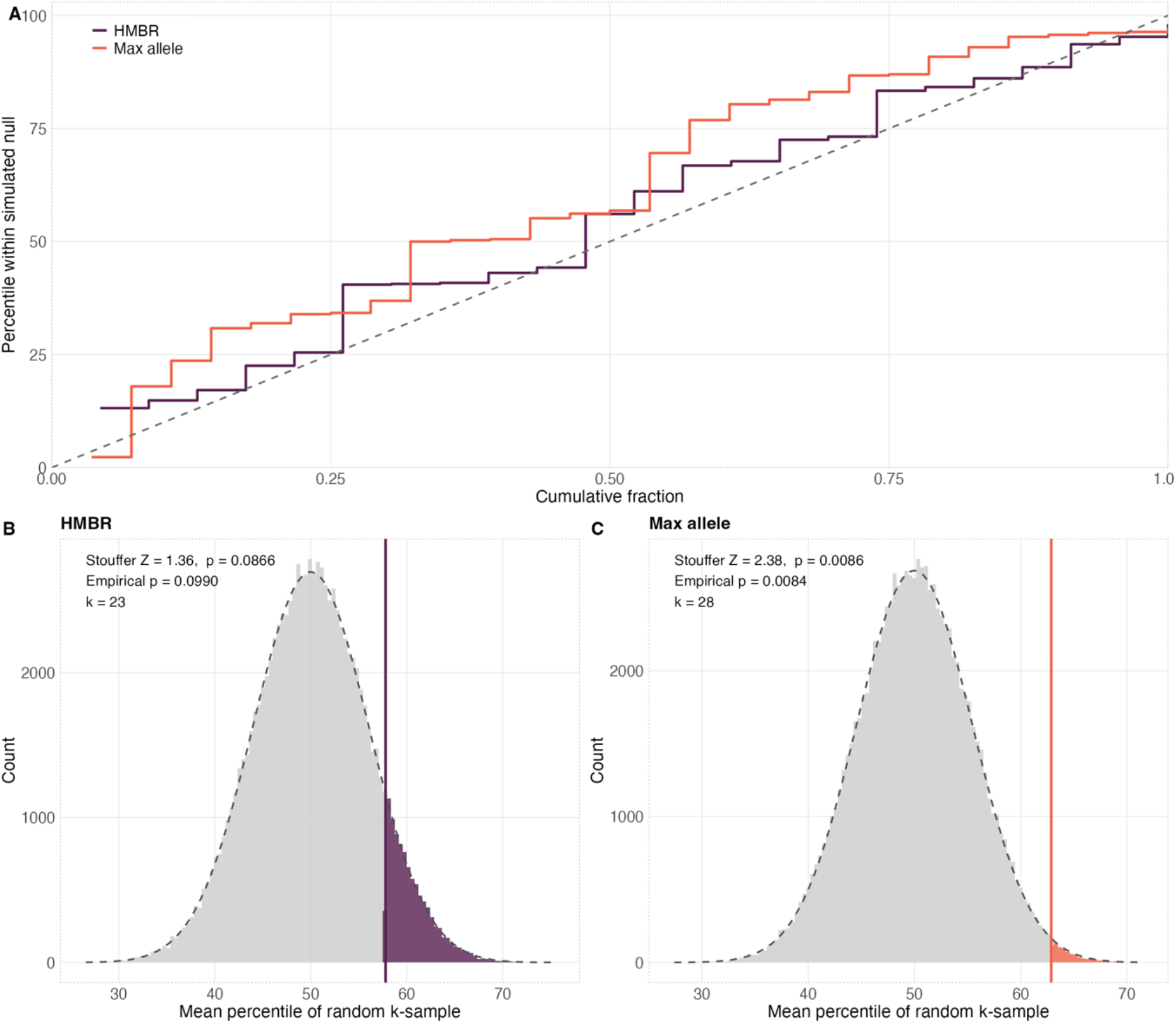
Statistical comparison of escape scores observed variants with an empirical score distribution. **A)** Empirical cumulative distribution function (ECDF) of percentile scores for observed variants under the harmonic mean best rank (HMBR, purple) and max-allele (orange) measures, relative to their respective simulated null distributions. A dashed diagonal line marks the expected ECDF under the null. Curves falling below the diagonal indicate a collective shift towards higher percentile scores. **B)** Distribution of mean percentile scores across 99,999 random draws of k = 23 variants from the HMBR simulated null. The observed mean percentile is shown as a vertical line; shaded bars indicate the proportion of draws meeting or exceeding the observed value (empirical p-value). A dashed curve shows the normal approximation to the null. **C)** As B, for the max-allele measure (k = 28).

## Discussion

We have here outlined CD8scape, a new software package facilitating the rapid analysis of potential CD8+ T cell escape mutations in viral genomes. Built around the netMHCpan platform, CD8scape allows the comparison of the potential effect of escape variants both within individual hosts, and in populations with known HLA compositions, potentially contributing to broader assessment of how virus evolution in immunocompromised hosts contributes to phenotypic change that affect the virus in a broader, population-level context. Applying our method to an example dataset, we have both replicated a previously published result and illustrated the potential for CD8scape to derive broader conclusions.

Our method incorporates two distinct metrics for evaluating CD8+ T cell escape. A simple maximum of the predicted escape scores for the different MHC alleles is perhaps the more natural of these two statistics when considering escape in a single patient, following a principle of immunodominance. However, when applied to a supertype panel, for which the frequency of individual haplotypes, rather than the frequency of their combinations, is generally known, the large number of potential alleles makes a simple maximum less representative of the whole. We present harmonic mean statistics as a proxy measure capturing an element of immunodominance across a broad set of alleles.

CD8scape is optimised for rapid calculations across panels and large numbers of peptides, with deduplication routines facilitating more efficient calculations than are achieved by the simple use of the original netMHCpan software.

## Code Availability

CD8scape is freely available under the GNU General Public License v3.0 at https://github.com/ewanwsmith/CD8scape.

## Acknowledgements

This work was supported by funding from the UK Medical Research Council (MC_UU_00034/1, MC_ST_00034, and MC_PC_ MR/Y002814/1).

## Author contributions

Conceptualization: ES; Data curation: ES, CI; Formal analysis: ES; Funding acquisition: CI, DR; Investigation: ES; Methodology: ES, CI; Project administration: CI; Resources: CI, DR; Software: ES; Supervision: JH, DR, CI; Validation: ES, CI; Visualization: ES; Writing – original draft: ES, CI; Writing – review & editing: ES, JH, DR, CI

---

**Figure S1.**
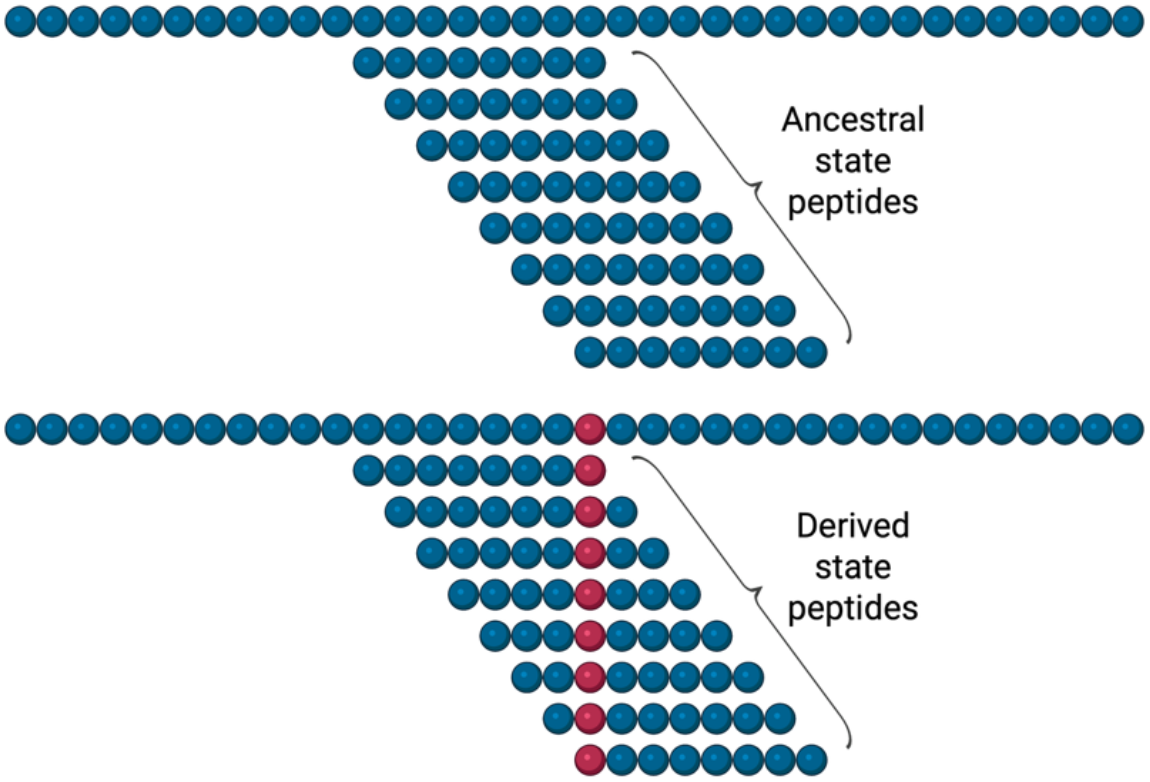
Schematic of CD8scape’s peptide generation strategy. For each variant locus, overlapping k-mers of lengths 8–11 are generated from both the ancestral and derived consensus sequences. Derived state peptides carry the variant amino acid (red). Created in BioRender.com.

**Figure S2.**
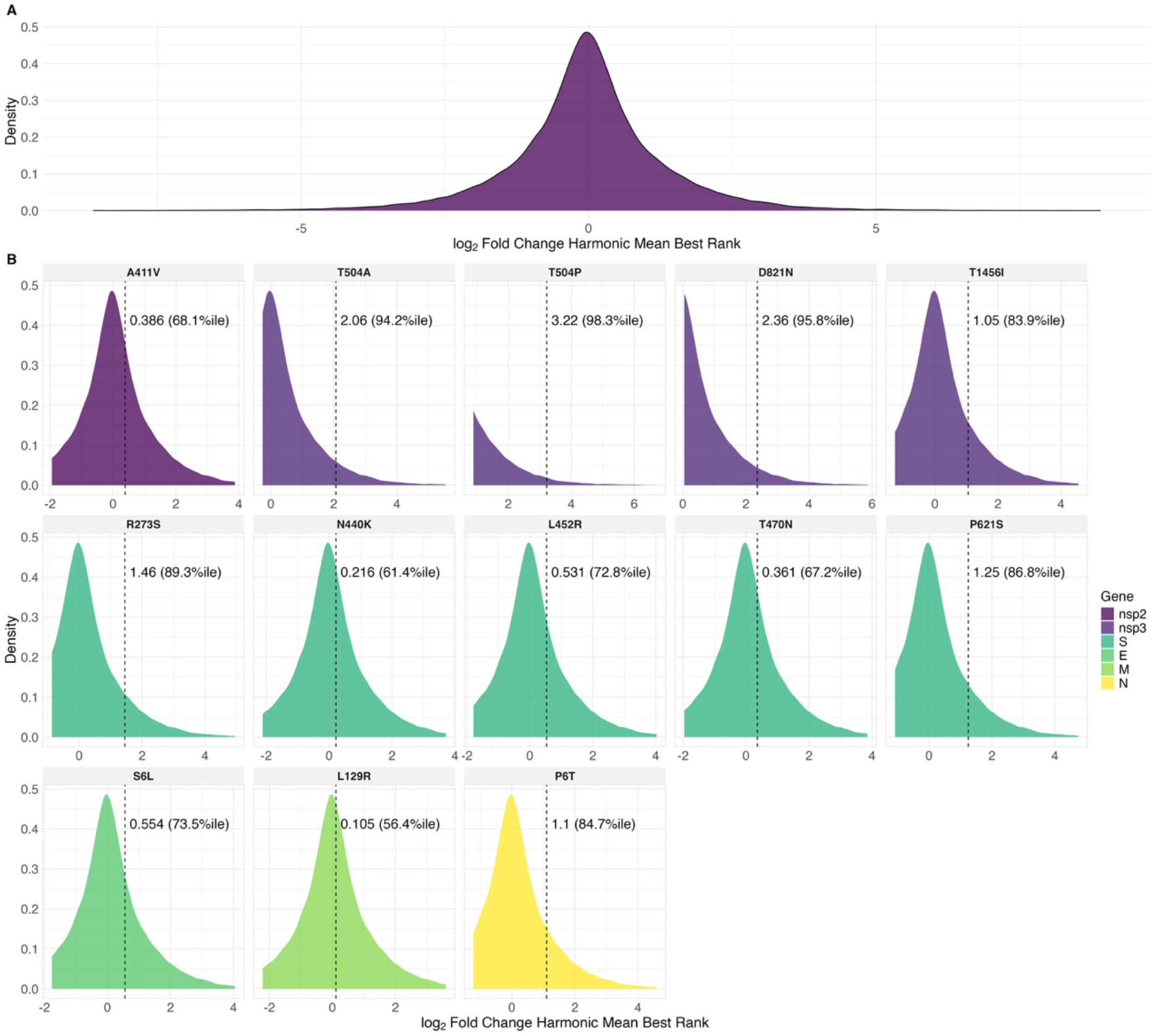
**A)** Empirical distribution of log_2_ fold-change in harmonic mean best EL rank across all possible non-synonymous mutations of the consensus SARS-CoV-2 genome. **B)** Percentile positions of observed variants with positive log_2_ fold-change on the simulated null distribution.

